# Skin commensal *Micrococcus luteus* Induces β-defensin-14 in keratinocytes and Enhances IL-17F Responses to Reduce *Candida auris* Colonization

**DOI:** 10.64898/2026.06.19.733356

**Authors:** Shrihari M Ganesh, Shankar Thangamani

## Abstract

*Candida auris* is a major multidrug-resistant fungal pathogen that predominantly colonizes human skin, leading to nosocomial transmission and outbreaks of systemic infections. Recent evidence suggests that *C. auris* co-colonizes with bacteria in the skin. However, the role of skin bacteria in the *C. auris* colonization is unclear. In this study, we investigated the role of *Micrococcus luteus*, a human skin commensal bacterium, on *C. auris* colonization of the skin. We identified that *M. luteus* pre-treatment in keratinocytes and mouse skin significantly reduces *C. auris* colonization. Mechanistically, we found that *M. luteus* induced β-defensin-14, a host antimicrobial peptide in skin keratinocytes, and enhanced IL-17F responses in T cells and innate lymphoid cells, thereby reducing *C. auris* skin colonization. These findings revealed potential microbiome-based therapeutics for the prevention and treatment of *C. auris* skin colonization and subsequent invasive infections in humans.

## Importance

*Candida auris* is an emerging fungal pathogen that poses a major threat to global public health. *C. auris* predominantly colonizes and persists in skin tissues. However, the host and microbiota factors that regulate *C. auris* skin colonization remain unclear. Understanding the interplay among the skin microbiota, the host, and *C. auris* is necessary to develop preventive and therapeutic approaches against *C. auris*. In this study, we identified that *Micrococcus luteus*, a skin commensal, reduces *C. auris* colonization by inducing host antimicrobial peptide and protective IL-17F response in the skin. These findings will open the door to modulating skin commensal bacteria to prevent and treat *C. auris* skin colonization and subsequent invasive fungal infections in humans.

### Observation

*Candida auris* is an emerging multidrug-resistant fungal pathogen. First identified in Japan in 2009, *C. auris* causes outbreaks of systemic infections and mortality among infected individuals [1–3]. *C. auris* has been identified as an urgent threat by the US Centers for Disease Prevention and Control (CDC) and is categorized as a critical priority group pathogen by the World Health Organization [4, 5]. The ability of *C. auris* to colonize the human skin leads to nosocomial transmission within health care facilities, resulting in major outbreaks [6, 7]. Recent studies suggest that *C. auris* coexists with bacteria and fungi on human skin [8–10]. A comparison of the distribution of skin microbiota between *C. auris*-positive and *C. auris-*negative individuals revealed that the skin of individuals colonized with *C. auris* had a significantly decreased abundance of beneficial commensal microbes [6]. However, the role of the commensal microbiota in *C. auris* skin colonization is poorly understood. In this study, we investigated the role of *Micrococcus luteus*, which is one of the major skin commensals in humans, on *C. auris* skin colonization [11–14]. Our findings suggest that *M. luteus* induces β-defensin-14, an antimicrobial peptide (AMP), in keratinocytes and enhances protective IL-17F responses, thereby reducing *C. auris* skin colonization. These results suggest that modulating beneficial skin microbiota could pave the way for the development of novel microbiome-based therapeutics to prevent and treat *C. auris* skin colonization in humans.

### *M. luteus* reduces *C. auris* colonization on keratinocytes by inducing β-defensin-14 expression through the aryl hydrocarbon receptor (AHR)

To investigate the role of *M. luteus* in *C. auris* skin keratinocytes, we pre-treated keratinocytes with live bacteria for 16 h, washed, and infected keratinocytes with *C. auris* (**Fig.1A**). *M. luteus* pre-treatment reduced *C. auris* adhesion and colonization in keratinocytes by ~ 3.9-fold compared to untreated cells (**Fig.1B**). Next, to confirm whether *in vitro* findings translate to *in vivo*, we used a mouse model of epicutaneous skin infection to determine if *M. luteus* application reduces *C. auris* colonization. *M. luteus* was topically applied to the mice’s skin for 5 days. Then, mice were infected with *C. auris* on day 6, and skin tissues were collected on day 9 to determine fungal burden (**Fig. 1C**). We observed a significant decrease in *C. auris* skin burden in *M. luteus* pretreated mice compared with PBS-treated mice (**Fig. 1D**). *M. luteus* pre-treated mice showed a 1.38 ± 0.24 log_10_ decrease in fungal load compared to PBS-treated mice. Because AMPs produced by keratinocytes play a critical role in host defense against skin pathogens [15, 16], we examined whether *M. luteus* induces AMPs in keratinocytes to reduce *C. auris* skin colonization. We examined the expression of major AMPs in keratinocytes. We found that the expressions of β-defensin-1 (*Defb1*) and β-defensin-3 (*Defb3*) were significantly downregulated, whereas β-defensin-4 (*Defb4*) and β-defensin-14 (*Defb14*) were significantly upregulated in keratinocytes pre-treated with *M. luteus* compared to untreated *C. auris*-infected cells (**Fig. 1E**). *Defb4* showed a ~26-fold increase, whereas *Defb14* showed a ~3-fold increase upon pre-treatment with *M. luteus*. Since β-defensins play a critical role in host defense against pathogens in the skin [17–19], we examined whether *M. luteus* reduces *C. auris* by inducing *Defb4* and *Defb14*. We used the siRNA approach to knock down these two defensins in keratinocytes. The efficacy of siRNA-mediated silencing of *Defb4* and *Defb14* was confirmed using qRT-PCR. As expected, a significant reduction in the mRNA levels of both AMPs was observed at 48h after transfection **(Fig. 1F**). Interestingly, we found that silencing of *Defb14* but not *Defb4* significantly increased fungal load compared to keratinocytes transfected with control siRNA **(Fig.1G**). Specifically, keratinocytes transfected with siDefb14 increased *C. auris* keratinocyte colonization by ~2.62-fold compared to that in negative control siRNA-transfected *M. luteus* pre-treated keratinocytes. These findings suggest that *M. luteus* reduces *C. auris* colonization of keratinocytes by upregulating *Defb14* expression. Skin commensals such as *Staphylococcus epidermidis* mediate keratinocyte host defense through the aryl hydrocarbon receptor (AHR) [20, 21], we aimed to determine whether *M. luteus* induces *Defb14* via the AHR pathway to reduce *C. auris* colonization. We found that *M. luteus* significantly increased AHR expression in keratinocytes’ *ex vivo* (**Fig.1H**). Next, we examined whether increased AHR functionally regulates *Defb14* expression and fungal colonization using CH223191, an AHR inhibitor. Keratinocytes treated with the CH223191 inhibitor showed significantly reduced *Defb14* expression compared with DMSO-treated controls (**Fig.1I**). CH223191 treatment reduced *Defb14* expression by approximately ~1.5-fold. Furthermore, treatment with CH223191 increased *C. auris* colonization 2-fold compared with DMSO-treated keratinocytes (**Fig. 1J**). Collectively, our results demonstrate that *M. luteus* reduces *C. auris* colonization of keratinocytes through AHR-dependent *Defb14* expression.

**Figure 1.**
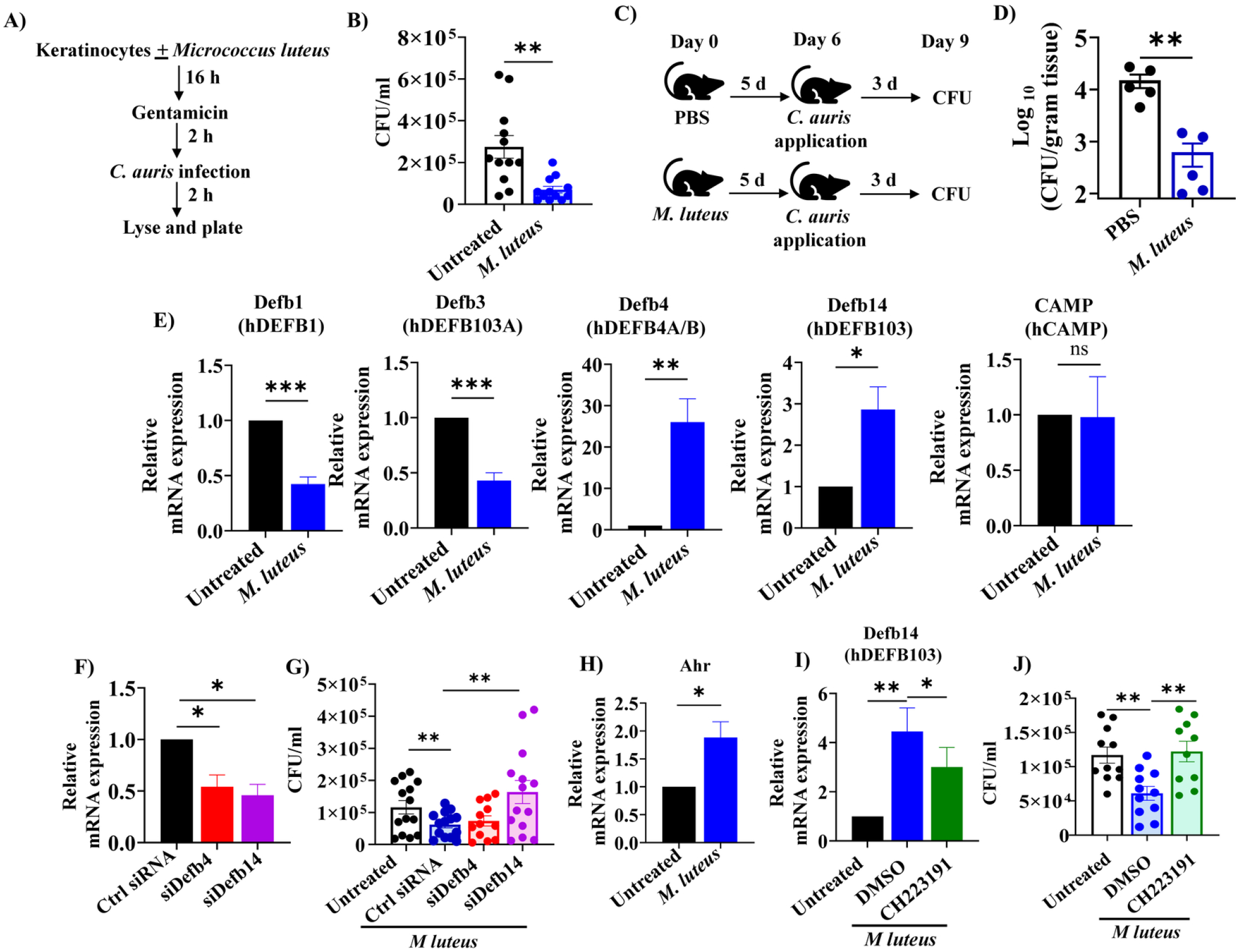
*M. luteus* induces *Defb14* through aryl hydrocarbon receptor (AHR) to reduce *C. auris* colonization in keratinocytes. **(A)** Schematic of experimental setup. **(B)** Keratinocytes pre-treated with live *M. luteus* and then infected with *C. auris*. Untreated groups used as controls. Keratinocytes were lysed and plated onto YPD plates to determine the fungal load. **(C)** Schematic of the mouse experimental setup. **(D)** Mouse skin was topically applied with PBS (or) live *M. luteus* for 5 days, then infected with *C. auris*. 72h post-*C. auris* infection, skin tissues were collected to determine the fungal load. **(E)** Keratinocytes pre-treated with or without live *M. luteus* and infected with *C. auris* to determine the AMPs expression by qRT-PCR. **(F)** Keratinocytes were transfected with the control siRNA, siDefb4, and siDefb14 to confirm the knockdown by qRT-PCR. **(G)** Keratinocytes transfected with the control siRNA, siDefb4, and siDefb14 were pre-treated with or without live *M. luteus* and infected with *C. auris* to determine the fungal load. **(H)** Keratinocytes pre-treated with or without live *M. luteus* and infected with *C. auris* to determine the AHR expression by qRT-PCR. **(I)** Keratinocytes pre-treated with or without live *M. luteus* in the presence or absence of AHR inhibitor and infected with *C. auris* to determine *Defb14* expression by qRT-PCR. **(J)** Keratinocytes pre-treated with or without live *M. luteus* in the presence or absence of AHR inhibitor and infected with *C. auris* to determine fungal load. Values are expressed as Mean ± SEM. Statistical significance was calculated using the Paired t-test (B, E-J) and Mann-Whitney U test (D). * *p* < *0*.*05*, ** p < 0.01, and *** *p* < 0.001 were considered significant.

### *M. luteus* induces IL-17F to restrict *C. auris* colonization in the skin

As IL-17-producing T cells and innate lymphoid cells (ILCs) play a crucial role in host defense against *C. auris* skin infection [2, 22], we investigated whether *M. luteus* regulates IL-17 immune response to control *C. auris* in the skin. We topically applied *M. luteus* to mice prior to *C. auris* skin infection, and after 14 days post-infection, skin tissues were collected to determine fungal burden and immune cell accumulation (**Fig. 2A**). We observed a significant decrease in *C. auris* skin burden in *M. luteus* pre-treated mice compared to PBS-treated mice, even 14 days post-infection (**Fig. 2B**). *M. luteus* pre-treated mice showed a 1.16 ± 0.22 log_10_ decrease in fungal load compared to PBS-treated mice. We examined T cell responses, including IL-17A, IL-17F, and IFNγ-producing CD4+, CD8+, and γδ+ T cells, as well as ILCs, in PBS and *M. luteus* pre-treated groups, as described previously [2]. We observed that the percentage and absolute numbers of CD4+ IL-17F+ T cells and IL-17F+ ILCs were significantly higher in *M. luteus* pre-treated mice than in PBS-treated mice (**Fig. 2C** and **2F**). We did not observe any significant differences in other cell types, including IFNγ+ T cells, between PBS and *M. luteus* pre-treated mice (**Fig. 2C-2F**). Next, we aim to determine whether *M. luteus*-induced IL-17F is necessary to protect against *C. auris* skin infection at 14 days post-infection. We depleted IL-17F-producing cells using an anti-mouse IL-17F antibody. Groups of mice were pre-treated with *M. luteus* and treated with (or) without anti-mouse IL-17F antibody, and the fungal burden was determined as indicated in Fig. 2G. Interestingly, mice pre-treated with *M. luteus* and treated with an IL-17F-neutralizing antibody showed a significant increase in fungal load compared to mice treated with an isotype control antibody (**Fig. 2H)**. Taken together, our findings indicate that the *M. luteus*-induced IL-17F response is critical for reducing *C. auris* skin colonization at 14 days post-infection.

**Figure 2.**
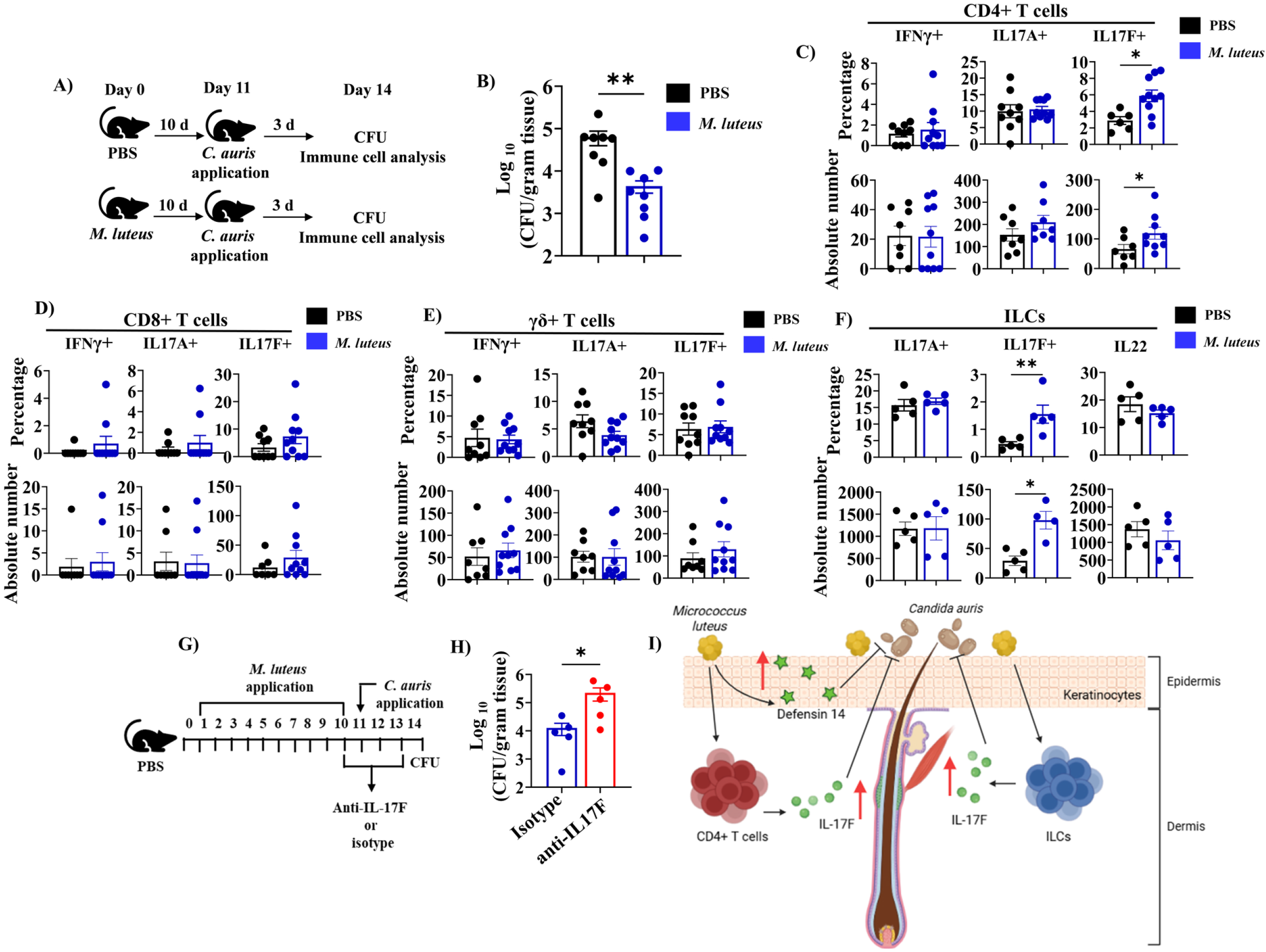
*M. luteus* enhances IL-17F response to reduce *C. auris* skin colonization. **(A)** Schematic of mouse experimental setup. **(B)** Mouse skin was topically applied with PBS (or) live *M. luteus* for 10 days, then infected with *C. auris*. 72h post *C. auris* infection, the skin tissues were collected to determine the fungal load. The skin tissues collected were used to determine: **(C)** The percentage and absolute numbers of CD4+ IFNγ+, CD4+ IL17A+, and CD4+ IL17F+ T cells, **(D)** Percentage and absolute numbers of CD8+ IFNγ+, CD8+ IL17A+, and CD8+ IL17F+ T cells, **(E)** Percentage and absolute numbers of γδ+ IFNγ+, γδ+ IL17A+, and γδ+ IL17F+ T cells, **(F)** Percentage and absolute numbers of IL17A+ ILCs, IL17F+ ILCs, and IL22+ ILCs. **(G)** Schematic of IL-17F neutralization experiment. **(H)** Mouse skin was topically applied with live *M. luteus* for 10 days, then infected with *C. auris*. 72h post *C. auris* infection, the skin tissues were collected to determine the fungal load. 100 μg/ml of isotype or anti-IL17F antibody was injected intraperitoneally on days 10-13. **(I)** Summary of how *M. luteus* reduces *C. auris* skin colonization. Figure generated using BioRender. Values are expressed as Mean ± SEM. Statistical significance was calculated using the Mann-Whitney U test. * *p* < 0.05 and ** *p* < 0.0, *** *p* < 0.001 were considered significant.

Collectively, our findings suggest that *M. luteus* induces *Defb14* host AMP in keratinocytes and enhances IL-17F-producing CD4+ T cells and ILCs responses, which together limit *C. auris* skin colonization at early and late time points post-infection (**Fig. 2I**). *M. luteus*, an understudied species, is among the major groups of skin commensals in humans that modulate host immune response in the skin [12, 23]. Recently, we identified that *C. auris* induces IFN-γ, which enhances fungal colonization in the skin [24]. Surprisingly, our findings in this study suggest that *M. luteus* induces AMPs and protective IL-17F responses without significantly altering IFN-γ response in the skin during *C. auris* infection. Given that the skin of *C. auris*-positive individuals lacks several commensal bacteria [6], understanding how commensal microbiota regulate *C. auris* will expand our knowledge of the microbiota and host immune factors that control *C. auris* skin colonization. Future studies to understand the molecular mechanisms, such as whether *M. luteus*, either directly or through secreted small molecules, induces AMPs, will open the door to harnessing the immunomodulatory properties of symbiotic factors from *M. luteus*, which could have therapeutic potential against *C. auris* [25]. This also paves the way for future interventions using *M. luteus* to treat *C. auris* skin colonization, prevent subsequent nosocomial transmission, and reduce mortality from invasive *C. auris* infection in humans [26].

## Materials and methods

### Cell culture

The mouse keratinocyte cell line (Kera308) was purchased from Cytion (USA) and maintained in Dulbecco’s Modified Eagle Medium (DMEM).

### Bacterial culture

The *M. luteus* strain SK58 used in this study was grown in Tryptic Soy Broth (TSB) medium overnight (16-18h) at 250rpm in a 37 °C shaker incubator and used for keratinocyte pretreatment or for treating mice with epicutaneous skin infection.

### Fungal culture

The *C. auris* AR0387 strain was cultured and grown in Yeast Peptone Dextrose (YPD) medium as described previously [24]

### Keratinocyte assay

Briefly, 1 × 10^5^ Kera308 cells/well were preincubated with *M. luteus* at a multiplicity of infection (MOI) of 1:10 for 16h. Then, the cells were washed 3 times with 1x PBS and incubated with gentamicin (50 μg/mL) for 2 h. Subsequently, cells were infected with *C. auris* at an MOI of 1:1 for 2h. After 2h, fungal load was determined by lysing cells with 1x PBS containing 0.02% Triton X-100 and plating onto YPD agar plates to determine the fungal load.

### Mice infection and fungal load determination

All animal experiments were performed according to the protocol described previously [2]. Animal studies and experimental protocols were approved by the Purdue University Institutional Animal Care and Use Committee (IACUC Protocol no. 2110002211). Briefly, 24h after dorsal skin hair removal, live *M. luteus* was applied epicutaneously to the shaved skin at a density of 1 × 10^7^ bacteria for 5 days or 10 days, and then 1 × 10^8^ *C. auris* was applied on day 6 or 11, respectively. Tissues were collected and homogenized to determine the fungal burden on day 9 or 14.

### Immune cell isolation and staining

Experiments were performed as described previously [2]. Briefly, skin tissues were minced and digested. Single-cell suspension was collected and activated for 4.5 hours at 37°C at 5% CO_2._ Cells were stained with the respective antibodies, acquired on an Attune NxT Flow Cytometer (Invitrogen, Carlsbad, CA, USA), and analyzed using FlowJo software (Eugene, OR, USA).

### Short interfering RNA (siRNA) experiment

For the siRNA experiment, 2 × 10^5^ kera308 cells/wells were transfected with control siRNA, siDefb14, and siDefb4 at 25nM for 48h. Following transfection, cells were washed with 1x PBS to determine the fungal load. For determining siRNA efficacy, 2 × 10^5^ kera308 cells were seeded in a 24-well plate for 24 hrs. Cells were transfected with control siRNA, siDefb14, and siDefb4 at 25nM for 48h. mRNA level expression was determined using RT-qPCR as described below

### RT-qPCR

To determine the mRNA expression levels of different AMPs, qRT-PCR was performed as described previously [24]. The following primers were used: *Defb1*, 5’-TCCTGGTGATGATATGTTTTCTTTTCT-3’and 5’-TGTTCTTCGTCCAAGACTTGTGA-3’; *Defb3*, 5’-GTCTCCACCTGCAGCTTTTAG-3’ and 5’-ACTGCCAATCTGACGAGTGTT-3’; *Defb4*, 5’-GCAGCCTTTACCCAAATTATC-3’ and 5’-ACAATTGCCAATCTGTCGAA-3’; *Defb14*, 5’-ATGAGGCTTCATTATCTGCTATTT-3’ and 5’-CTACTTCTTCTTTCGGCAGCATTT-3’; *Camp*, 5’-GCTGATTCTTTTGACATCAGCTGTAA-3’ and 5’-GCCAGCCGGGAAATTTTCT-3’

### Statistical analysis

Statistical analysis was performed using the GraphPad Prism 9.4.1 (GraphPad Software, La Jolla, CA, USA). *P* values ≤ 0.05 were considered significant.

## References

1. Lionakis, M.S. and A. Chowdhary, Candida auris Infections. N Engl J Med, 2024. 391(20): p. 1924–1935.

2. Datta, A., et al., Differential skin immune responses in mice intradermally infected with Candida auris and Candida albicans. Microbiol Spectr, 2023. 11(6): p. e0221523.

3. Santana, D.J., et al., A Candida auris-specific adhesin, Scf1, governs surface association, colonization, and virulence. Science, 2023. 381(6665): p. 1461–1467.

4. Kadri, S.S., Key Takeaways From the U.S. CDC’s 2019 Antibiotic Resistance Threats Report for Frontline Providers. Crit Care Med, 2020. 48(7): p. 939–945.

5. Organization, W.H., WHO fungal priority pathogens list to guide research, development and public health action. World Health Organization, 2022: p. 48.

6. Proctor, D.M., et al., Integrated genomic, epidemiologic investigation of Candida auris skin colonization in a skilled nursing facility. Nat Med, 2021. 27(8): p. 1401–1409.

7. Dire, O., et al., Survival of Candida auris on environmental surface materials and low-level resistance to disinfectant. J Hosp Infect, 2023. 137: p. 17–23.

8. Sansom, S.E., et al., Rapid Environmental Contamination With Candida auris and Multidrug-Resistant Bacterial Pathogens Near Colonized Patients. Clin Infect Dis, 2024. 78(5): p. 1276–1284.

9. Proctor, D.M., et al., Integrated genomic, epidemiologic investigation of Candida auris skin colonization in a skilled nursing facility. Nat Med, 2021. 27(8): p. 1401–1409.

10. Proctor, D.M., et al., Clonal Candida auris and ESKAPE pathogens on the skin of residents of nursing homes. Nature, 2025. 639(8056): p. 1016–1023.

11. McLoughlin, I.J., et al., Skin Microbiome-The Next Frontier for Probiotic Intervention. Probiotics Antimicrob Proteins, 2022. 14(4): p. 630–647.

12. Jain, R., et al., Efficacy of a topical live probiotic in improving skin health. Int J Cosmet Sci, 2025. 47(3): p. 488–496.

13. Chiller, K., B.A. Selkin, and G.J. Murakawa, Skin microflora and bacterial infections of the skin. J Investig Dermatol Symp Proc, 2001. 6(3): p. 170–4.

14. Heo, Y.M., et al., Skin benefits of postbiotics derived from Micrococcus luteus derived from human skin: an untapped potential for dermatological health. Genes Genomics, 2024. 46(1): p. 13–25.

15. Bier, K. and B. Schittek, Beneficial effects of coagulase-negative Staphylococci on Staphylococcus aureus skin colonization. Exp Dermatol, 2021. 30(10): p. 1442–1452.

16. Parlet, C.P., M.M. Brown, and A.R. Horswill, Commensal Staphylococci Influence Staphylococcus aureus Skin Colonization and Disease. Trends Microbiol, 2019. 27(6): p. 497–507.

17. Dinulos, J.G., et al., Keratinocyte expression of human beta defensin 2 following bacterial infection: role in cutaneous host defense. Clin Diagn Lab Immunol, 2003. 10(1): p. 161–6.

18. Dong, X., et al., Keratinocyte-derived defensins activate neutrophil-specific receptors Mrgpra2a/b to prevent skin dysbiosis and bacterial infection. Immunity, 2022. 55(9): p. 1645–1662 e7.

19. Niyonsaba, F., et al., Antimicrobial peptides human beta-defensins stimulate epidermal keratinocyte migration, proliferation and production of proinflammatory cytokines and chemokines. J Invest Dermatol, 2007. 127(3): p. 594–604.

20. Stockinger, B., et al., The aryl hydrocarbon receptor: multitasking in the immune system. Annu Rev Immunol, 2014. 32: p. 403–32.

21. Rademacher, F., et al., Staphylococcus epidermidis Activates Aryl Hydrocarbon Receptor Signaling in Human Keratinocytes: Implications for Cutaneous Defense. J Innate Immun, 2019. 11(2): p. 125–135.

22. Huang, X., et al., Murine model of colonization with fungal pathogen Candida auris to explore skin tropism, host risk factors and therapeutic strategies. Cell Host Microbe, 2021. 29(2): p. 210–221 e6.

23. Elias, A.E., et al., A skin isolate of Micrococcus luteus negates the Staphylococcus aureus-induced release of type 2 cytokines from keratinocytes. Front Immunol, 2026. 17: p. 1711723.

24. Das, D., et al., The Emerging Fungal Pathogen Candida auris Induces IFNgamma to Colonize the Skin. PLoS Pathog, 2025. 21(4): p. e1013114.

25. Severn, M.M., et al., The Ubiquitous Human Skin Commensal Staphylococcus hominis Protects against Opportunistic Pathogens. mBio, 2022. 13(3): p. e0093022.

26. Zhao, G., et al., Adhesin Als4112 promotes Candida auris skin colonization through interactions with keratinocytes and extracellular matrix proteins. Nat Commun, 2025. 16(1): p. 5673.

